# “Metabolic contest”, a new way to control carbon source preference

**DOI:** 10.1101/800839

**Authors:** Stefan Allmann, Marion Wargnies, Edern Cahoreau, Marc Biran, Nicolas Plazolles, Pauline Morand, Erika Pineda, Hanna Kulyk, Corinne Asencio, Oriana Villafraz, Loïc Rivière, Emmanuel Tétaud, Brice Rotureau, Arnaud Mourier, Jean-Charles Portais, Frédéric Bringaud

## Abstract

Microorganisms must make the right choice for nutrient consumption to adapt to their changing environment. As a consequence, bacteria and yeasts have developed regulatory mechanisms involving nutrient sensing and signaling, allowing to redirect cell metabolism to maximize the consumption of an energy-efficient carbon source. Here, we report a new mechanism, named “metabolic contest”, for regulating the use of carbon sources without nutrient sensing and signaling. In contrast to most microorganisms, trypanosomes show a glycerol-to-glucose preference that depends on the combination of three conditions: (*i*) the sequestration of both metabolic pathways in the same subcellular compartment, here in the peroxisomal-like organelles named glycosomes; (*ii*) the competition for the same substrate, here ATP, with the first enzymatic step of the glycerol and glucose metabolic pathways being both ATP-dependent (glycerol kinase and hexokinase, respectively) and (*iii*) an unbalanced activity between the competing enzymes, here the glycerol kinase activity being ~80-fold higher than the hexokinase activity.

## INTRODUCTION

Microorganisms show a remarkable ability to optimize the utilization of resources occurring in their environment. When various nutrients are available, microbes are able to use them in a sequential order, according to the efficiency by which they can metabolize these different compounds. The result is a hierarchical utilization of nutrients, which is microorganism- and strain-specific. The sequential utilization of nutrients is due to catabolic repression, a phenomenon by which the expression of enzymes involved in the catabolism of a compound are repressed in the presence of a more efficient nutrient. Not only this allows the optimization of resource utilization, but also it avoids energy spillage by preventing the biosynthesis of unnecessary proteins. Catabolite repression was first described in bacteria with the discovery of diauxie, i.e. the occurrence of two distinct growth phases due to the sequential utilization of two carbon sources (generally sugars), and is also observed in yeasts and fungi. The molecular mechanisms underlying catabolite repression were detailed after the first description in 1964 of the phosphoenolpyruvate:sugar phosphotransferase system (PTS) (Kundig et al., 1964). This is a carbohydrate transport and phosphorylation system composed of three protein complexes that regulates numerous cellular processes by either phosphorylating its target proteins or by interacting with them in a phosphorylation-dependent manner (Galinier and Deutscher, 2017). Indeed, catabolite repression relies on the presence and activity of specific sensing and signaling proteins repressing the expression of genes encoding transport systems or catabolic enzymes (Gorke and Stulke, 2008; Kayikci and Nielsen, 2015; Lengeler, 2015).

*Trypanosoma brucei* is a unicellular eukaryote belonging to the Kinetoplastid order, that causes Human African Trypanosomiasis, also known as sleeping sickness (Buscher et al., 2017). The transmission of these parasites is ensured by a haematophagous insect vector of the genus *Glossina*, also called tsetse. During their complex life cycle within two different hosts, trypanosome parasites face highly diverse micro-environments and nutritional conditions, urging the need for highly efficient metabolic adaptation mechanisms. Here, we report a novel molecular mechanism for the management of available resources, named “metabolic contest”, illustrated in the procyclic forms (PCF) of trypanosomes present in the digestive tract of the insect vector. PCF prefer glycerol, a gluconeogenic carbon source, to glucose. The glycerol preference is due to a competition for ATP between glycerol kinase and hexokinase, which is strongly in favor of the first enzyme in the presence of glycerol and prevents glucose utilization until glycerol is totally consumed. The subcellular localization of the two competing enzymes into the glycosome (Figure 1A), a specialized organelle hosting the first reactions of the glycolytic pathway in these parasites (Opperdoes and Borst, 1977), is a key feature for such a process.

**Figure 1.**
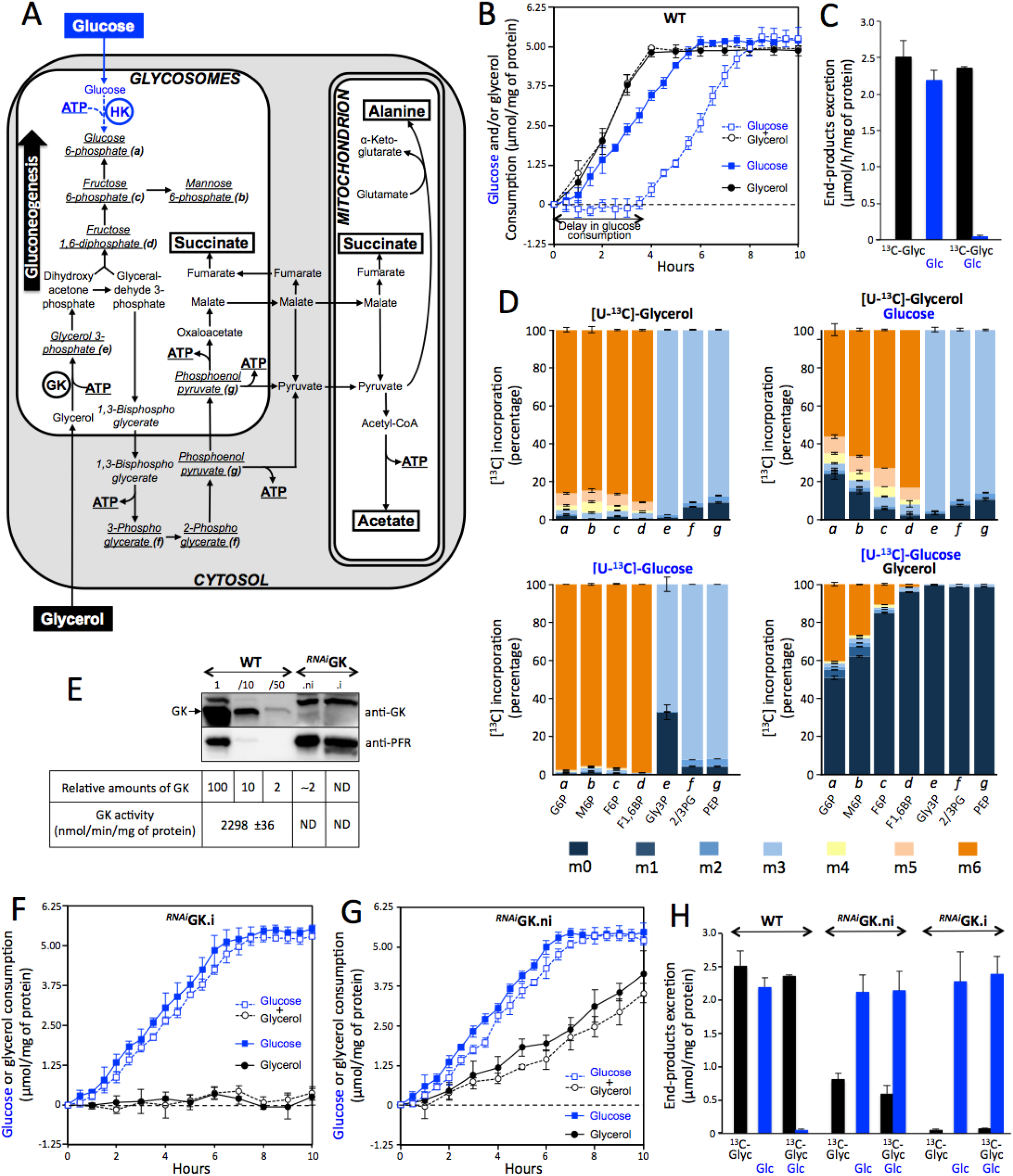
Procyclic trypanosomes prefer glycerol to glucose. (A) Schematic representation of glycerol (black) and glucose (blue) metabolisms in PCF trypanosomes. The metabolic end-products are shown in a rectangle and metabolites analyzed by IC-HRMS are underlined and in italic. The ATP molecules consumed and produced by substrate level phosphorylation are shown as well as the enzymes hexokinase (HK) and glycerol kinase (GK). (B) Glucose and glycerol consumptions by PCF trypanosomes incubated in glucose (2 mM), glycerol (2 mM) and glucose + glycerol (2 mM each) conditions. (C) Metabolic end-products produced by PCF trypanosomes from [U-^13^C]-glycerol (^13^C-Glyc) and/or glucose (Glc), as measured by ^1^H-NMR spectroscopy (the values are calculated from the data presented in Table S1). (D) IC-HRMS analysis of intracellular metabolites collected from PCF trypanosomes after incubation with 2 mM [U-^13^C]-labeled carbon sources in the presence or not of unlabeled carbon sources, as indicated on the top of each graph. The figure shows the proportion (%) of molecules having incorporated 0 to 6 ^13^C atoms (m0 to m6, color code indicated below). 2/3PG means 2- or 3-phosphoglycerate since these two metabolites are undistinguished by IC-HRMS. *: in this incubation condition the absolute amount of F1,6BP was very low and only the m6 isotopomer could be detected. (E) Western blot analysis of total protein extracts from the parental (WT) and tetracycline-induced (.i) or uninduced (.ni)^*RNAi*^ GK cell lines probed with anti-GK and anti-PFR (paraflagellar rod) immune sera. The table below the blots shows the relative levels of GK expression in 5 × 10^6^(1), 5 × 10^5^ (/10) and 10^5^ (/50) parental cells and 5 × 10^*6RNAi*^ GK.ni and ^*RNAi*^GK.i cells, as well as the corresponding GK activities (ND, not detectable). (F-G) Glucose and glycerol consumption by the ^*RNAi*^GK.i (F) ^*RNAi*^GK.ni or (G) ^*RNAi*^GK mutant cell line incubated in glucose (2 mM), glycerol (2 mM) and glucose + glycerol (2 mM each) conditions. (H) Production of metabolic end-products by the parental (WT), ^*RNAi*^GK.ni and ^*RNAi*^GK.i cell lines from [U-^13^C]-glycerol (^13^C-Glyc) and/or glucose (Glc), as measured by ^1^H-NMR spectroscopy (the values are calculated from the data presented in Table S1).

## RESULTS AND DISCUSSION

### Glycerol down-regulates glucose catabolism

We first measured the consumption of glucose or glycerol by procyclic forms (PCF) of *T. brucei* parasites maintained in culture in SDM79 medium supplemented with either glycerol or glucose or both. Glycerol was consumed by PCF faster than glucose whatever the conditions. The rate of glycerol consumption was not affected by the presence of glucose. In contrast, the latter was not consumed as long as glycerol was present in the medium (Figure 1B). After glycerol exhaustion, glucose was consumed at a rate similar to glucose alone conditions. This absence of glucose consumption in the presence of glycerol clearly showed a strong catabolic repression-like effect exerted by glycerol on glucose. It is noteworthy that the consumption of glucose started as soon as glycerol is exhausted, with no delay. Consistently, the growth profile of PCF trypanosomes in glucose + glycerol conditions was not characteristic of a diauxic effect (Figure S1), i.e. there was a single growth phase not two, in contrast to observations made in yeast when several carbon sources are available, the first one consumed (glucose) being the one that ensures the highest growth rate (Galdieri et al., 2010).

To confirm this glycerol preference in PCF trypanosomes, we have monitored the metabolic fate of ^13^C-labeled glycerol ([U-^13^C]-glycerol) alone or in combination with equimolar amounts of unlabeled glucose. The analysis of metabolic end-products (Figure 1C and Table S1) by ^1^H-NMR spectroscopy (Bringaud et al., 2015) allowed determining the respective contribution of [U-^13^C]-glycerol (labeled compounds) and glucose (unlabeled compounds). Trypanosomes mostly excreted acetate and succinate from glycerol or glucose (see Figure 1A). The rate by which glycerol is converted into these compounds was only slightly modified by the presence of glucose. In contrast, the conversion of glucose into acetate and succinate was reduced by ~50-fold in the presence of [U-^13^C]-glycerol. Moreover, the small production of lactate and alanine from glucose observed in the absence of glycerol was abolished in its presence. This significant reduction in the conversion of glucose into end-products in the presence of glycerol was correlated with the 20.4-fold decrease in glucose consumption in the same conditions, confirming that glucose metabolism was strongly down-regulated in the presence of glycerol.

Since production of glucose 6-phosphate (G6P) through gluconeogenesis is essential in the absence of glucose (see Figure 1A), we measured the incorporation of ^13^C-label into glycolytic intermediates by IC-HRMS in PCF trypanosomes incubated with [U-^13^C]-glycerol. In this experiment, most hexose-phosphate glycolytic intermediates were fully [^13^C]-labeled (88.2% ±1.8 of total molecules on average) after 2 hours of incubation with [U-^13^C]-glycerol as the sole carbon source (Figure 1D, top left). Addition of equal amounts of unlabeled glucose only slightly reduced ^13^C-incorporation into hexose-phosphates, with an average of 69.9% ±9.7 fully [^13^C]-labeled molecules (Figure 1D, top right). To confirm this glycerol-to-glucose preference, the equivalent experiment was performed with [U-^13^C]-glucose (Figure 1D, bottom). Addition of equal amounts of unlabeled glycerol abolished incorporation of ^13^C from [U-^13^C]-glucose into triose-phosphates and fructose 1,6-bisphosphate (F1,6BP). The ^13^C-incorporation into G6P was strongly reduced (40% ±0.9 vs 98% ±0.1 fully [^13^C]-labeled molecules in the presence and absence of glycerol, respectively).

Altogether, these data demonstrate that PCF trypanosomes significantly prefer glycerol to glucose for the production of hexose phosphates, including the first glycolytic intermediate, *i.e*. G6P. These data also suggest that the hexokinase (HK), producing G6P from glucose and/or some glucose transporters may be the target(s) of the glycerol-induced down-regulation of glucose metabolism. As far as we are aware, PCF trypanosomes are the only extracellular microorganisms described to date showing a glycerol-to-glucose preference. *T. brucei* PCF are also the only known glycolytic-competent lower eukaryotes performing gluconeogenesis in the presence of glucose.

### Glycerol metabolism is critical for the reduction of glucose catabolism

To further study glycerol metabolism in PCF trypanosomes, the expression of the first enzyme of the glycerol pathway was down-regulated by a RNAi silencing approach simultaneously targeting the five tandemly arranged *GK* genes (Tb927.9.12550 - Tb927.9.12630) under control of an tetracycline-inducible system. In the absence of tetracycline, the uninduced ^*RNAi*^GK cell line (^*RNAi*^GK.ni) presented a strong constitutive leakage of the RNAi silencing system with a 50-fold reduction of the GK protein content, reducing the overall GK enzyme activity to undetectable level (Figure 1E). The residual GK protein level could be further reduced after tetracycline induction (^*RNAi*^GK.i). Therefore, the direct involvement of GK in glycerol metabolism of these cells was determined by measuring glycerol consumption and release of metabolic end-products in glycerol conditions (Figure 1F-H and Table S1). Both the glycerol consumption and acetate/succinate production from glycerol metabolism were almost abolished in the ^*RNAi*^GK.i mutant, demonstrating that there is no alternative to GK for glycerol breakdown in PCF. Interestingly, the presence of glycerol did not affect glucose consumption by the ^*RNAi*^GK.i mutant (Figure 1F), indicating that the presence of glycerol in the medium is not *per se* responsible of glucose repression. In other words, glycerol does not directly affect glucose uptake and metabolism, which implies that intracellular glycerol metabolism is required to repress glucose metabolism. It is also important to mention that replacing glucose by glycerol did not affect growth of the ^*RNAi*^GK.i mutant, given that proline was the main carbon source used in these conditions as in the insect vector midgut.

The analysis of the ^*RNAi*^GK.ni cell line also provided relevant information regarding the unexpected role of GK activity in the glycerol-to-glucose preference process. First, the consumption of glycerol (Figure 1G) and its conversion into end-products (Figure 1H and Table S1) were reduced only by 3.5-fold and 3.1-fold, respectively, in the ^*RNAi*^GK.ni mutant as compared to the parental cells, while the GK expression was ~50-fold down-regulated (Figure 1E). One can extrapolate from these data that a reduction of GK activity by at least 90% would not affect the glycerol metabolism flux, which would further highlight a large excess of GK activity in PCF (in the range of 10-fold). Second, glucose metabolism was no longer repressed by glycerol in the ^*RNAi*^GK.ni cells, that consumed glucose at the same rate as the parental cells, without any glycerol-induced delay, although the glycerol consumption remained constant over the 10 hours of incubation (Figure 1G). This strongly suggests that the abolition of the glycerol-to-glucose preference in the ^*RNAi*^GK.ni cells could be the consequence of the 50-fold reduction of the GK activity.

### The glucose catabolism repression is due to a large excess of GK activity

Interestingly, GK activity was approximately 80-fold higher than HK activity in total PCF extracts (Figure 2B), using the enzymatic assays described in Figure 2A. Since these two glycosomal enzymes are competing for the same ATP pool (glycosomal), we hypothesized that this significant difference of activity would favor glycerol metabolism and disfavor glucose metabolism, and would hence explain the unique repression of glucose by glycerol. To test this hypothesis, we measured HK activity in the presence or absence of glycerol under incubation conditions compatible with HK and GK activities. Importantly, the enzymatic assay included 0.6 mM ATP, which corresponds to the measured glycosomal concentration (Bakker et al., 1997). The presence of glycerol in the assay induced a 15-fold reduction of HK activity in the parental cell extracts, but not in the ^*RNAi*^GK.ni and ^*RNAi*^GK.i cell extracts (Figure 2B), which demonstrates that the conversion of glycerol into Gly3P, but not the presence of glycerol *per se*, inhibits HK activity and thus glucose metabolism. In contrast, GK activity was not impaired by the presence of glucose in the parental cell line extract (Figure 2B).

**Figure 2.**
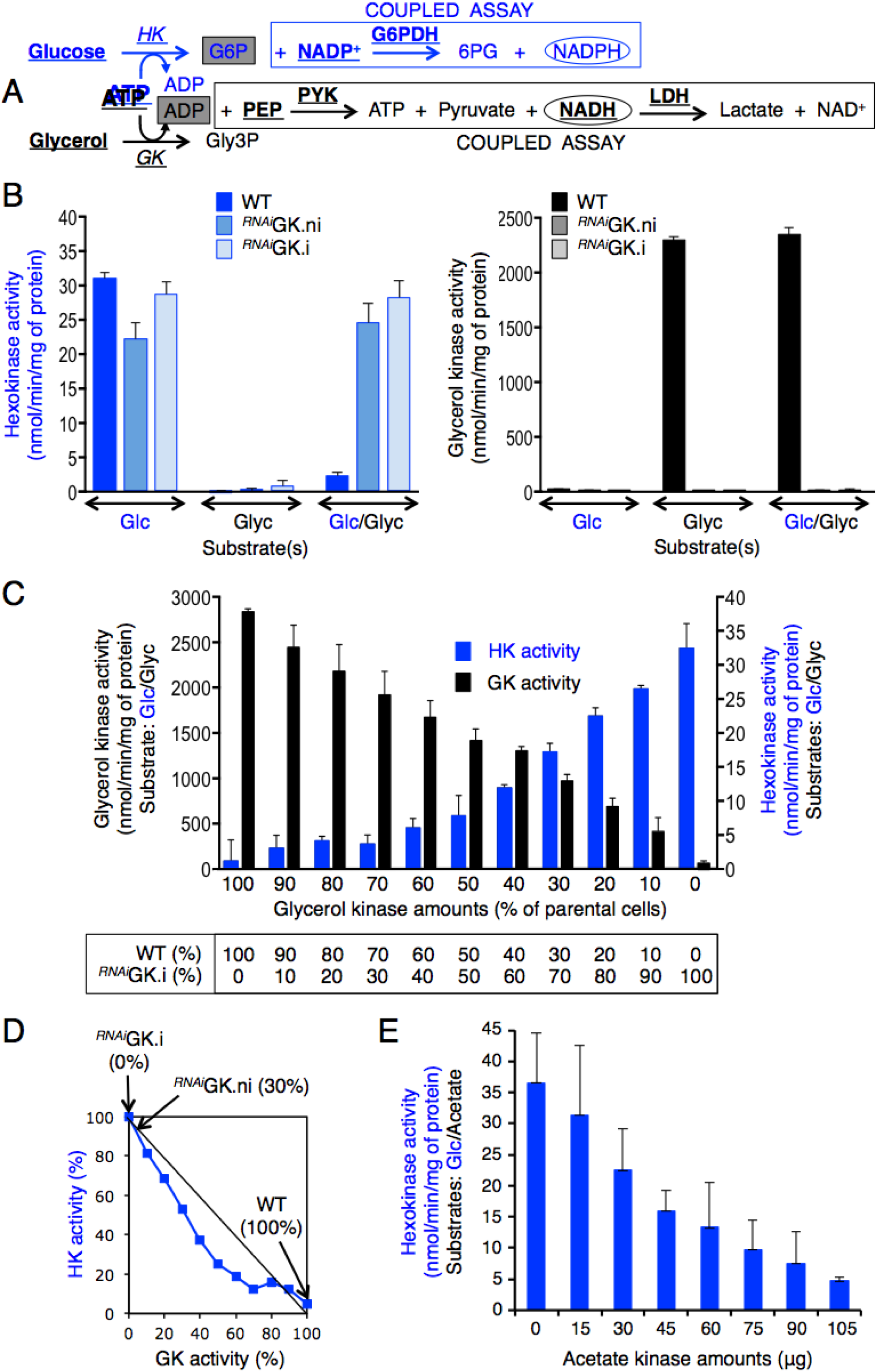
The glycerol-preference is the consequence of the high excess of GK activity. (A) Enzymatic assays used for the quantification of HK and GK activities. The bold and underlined substrates and enzymes are included in the assay for production of NADPH (HK assay) and consumption of NADH (GK assay) that are detected by spectrometry at 340 nm. (B) HK (left panel) and GK (right panel) activities in total cell extracts (WT, ^*RNAi*^GK.i and ^*RNAi*^GK.ni) determined in the presence of glucose (Glc), glycerol (Glyc) or equal amounts of glucose and glycerol (Glc/Glyc). (C) GK and HK activities in different combinations (indicated in the table below the graph) of total cell extracts from the parental (WT) and the ^*RNAi*^GK.i cell lines. The amounts of HK remain the same in all samples, while the amounts of GK present in the parental samples are diluted with the GK-depleted ^*RNAi*^GK.i samples. The HK and GK activities were determined in the presence of both glucose and glycerol, as performed in the Glc/Glyc conditions (see B). (D) Expression of HK activity as a function of GK activity. The values into brackets means the rate of glycerol consumption in the ^*RNAi*^GK.ni and ^*RNAi*^GK.i cells compared to the parental cell (100%). (E) HK activity in the presence of 10 mM acetate and increasing amounts of acetate kinase.

We took advantage of the fact that both the parental and ^*RNAi*^GK.i cell extracts were displaying similar HK activities (see Figure 2B) to further characterize the GK-derived inhibition effect on HK activity as a function of the HK/GK activity ratio, by diluting the parental cell extract with different volumes of the ^*RNAi*^GK.i cell extract (Figure 2C). As expected, HK activity was equivalent in all the samples in the presence of 10 mM glucose (data not shown). However, the addition of 10 mM glycerol decreased HK activity and this effect was dependent on the HK/GK ratio. Indeed, a reverse correlation between HK and GK activities was observed (Figure 2D), which was consistent with our hypothesis that both enzymes are competing for the same ATP pool. To confirm that this inhibitory effect was due to GK activity rather than to any other activities or biochemical properties of the enzyme, GK and glycerol were replaced by an acetate kinase and acetate in the same HK activity assay. As anticipated, HK activity was inhibited concomitantly to increasing amounts of acetate kinase (Figure 2E).

Glycerol metabolism was also investigated in the PCF of *T. congolense*, a trypanosome species closely related to *T. brucei*, bearing a single GK copy, instead of 5 copies in the *T. brucei* genome. GK activity was 5.5-fold lower in *T. congolense* compared to *T. brucei*, with a GK/HK activity ratio ~20-fold lower in *T. congolense* (Figure S2). Glycerol metabolism did not impair glucose consumption in *T. congolense* as shown by (*i*) the persistence of high HK activity in the presence of glycerol and (*ii*) the simultaneous consumption of glucose and glycerol (Figure S2), as observed for the *T. brucei* ^*RNAi*^GK.ni cell line (Figure 1G). In addition, we previously reported that the bloodstream forms (BSF) of *T. brucei* show no preference for glycerol or glucose (Pineda et al., 2018). This is consistent with the unaffected HK activity in the presence of glycerol and with the 28-fold increase of HK activity in BSF as compared to PCF, while GK activities are equivalent in the two forms of the parasites (Figure S2). Overall, our hypothesis is confirmed by these last data showing a strong correlation between the glycerol-to-glucose preference and the large excess of GK activity compared to HK activity.

### Accumulation of Gly3P and F1,6BP does not inhibit HK activity

To investigate whether the parasite metabolic profiles were dependent on the growth conditions, we determined their absolute intracellular concentrations by IC-HRMS by adding an internal standard ([U-^13^C]-labeled *E. coli* extract) into the *T. brucei* cell extracts, as described before (Ebikeme et al., 2010). Among the glycolytic intermediates analyzed, F1,6BP and glycerol 3-phosphate (Gly3P) accumulated approximately 5-fold and 49-fold more, respectively, in parental cells grown on glycerol as compared to those grown on glucose (Figure 3A). This significant Gly3P accumulation was confirmed by using an enzymatic determination (Figure 3B). It is noteworthy that this Gly3P accumulation persisted in the parental cells incubated with equal amounts of glycerol and glucose, while it was abolished for the ^*RNAi*^GK.ni mutant (Figure 3B). Consequently, one may consider that accumulation of intracellular amounts of Gly3P would inhibit the HK and prevents G6P production from glucose. To test this hypothesis, HK activity was measured in the presence of Gly3P in ^*RNAi*^GK.i mutant extracts, rather than parental cells, in order to prevent any interferences of the glycerol metabolism on the assay. Addition of up to 40 mM Gly3P did not significantly affect the *in vitro* HK activity (Figure 3C), which is consistent with previously published data (Dodson et al., 2011). It is also noteworthy that HK activity was not inhibited by adding up to 5 mM F1,6BP neither (Figure 3D). This shows that the accumulation of G3P or F1,6BP is not responsible for the glycerol-to-glucose preference.

**Figure 3.**
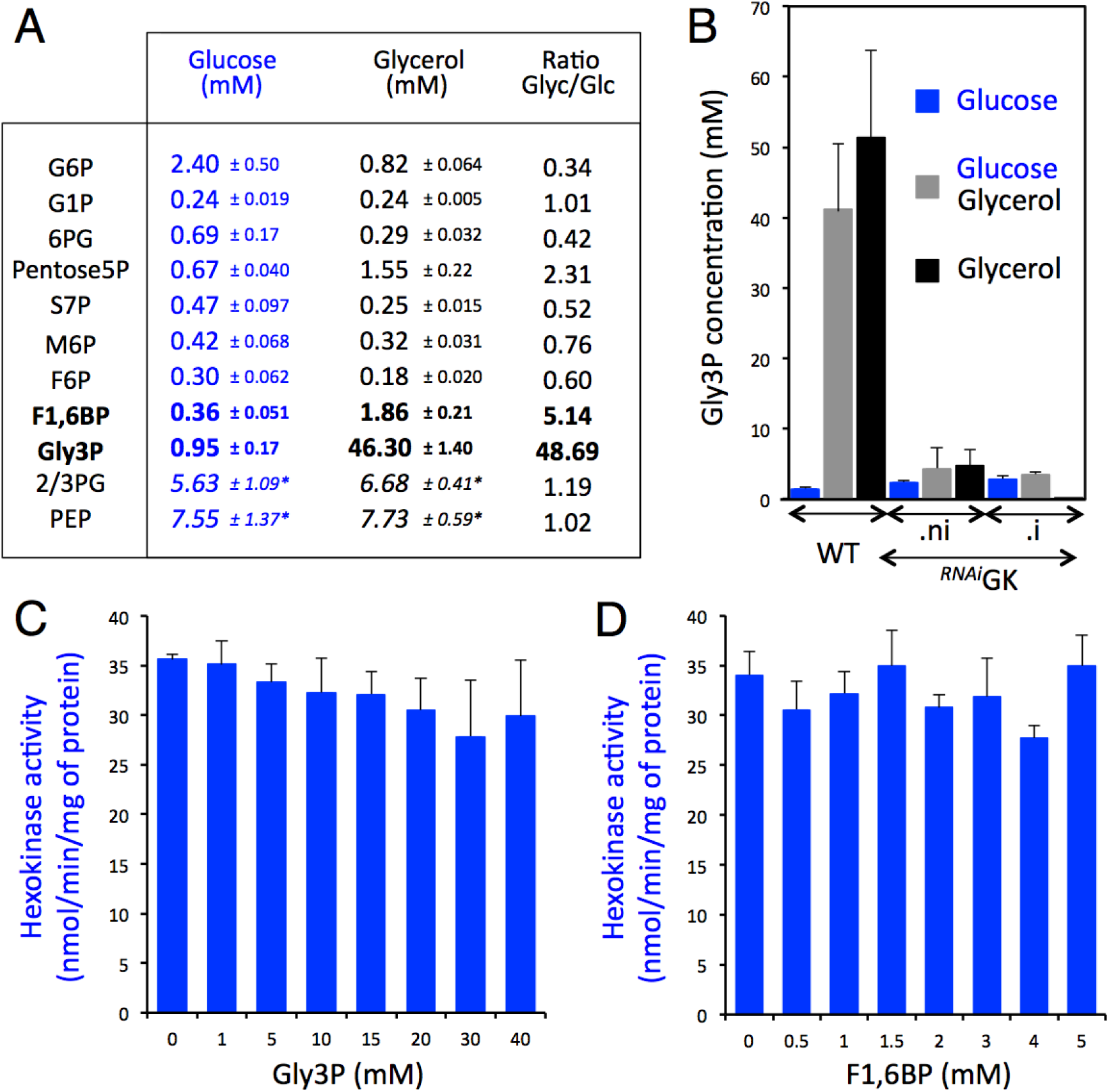
Analysis of intracellular metabolite concentrations. (A) Intracellular concentrations of metabolites in PCF trypanosomes grown in the presence of 10 mM glucose or glycerol. The concentrations of the 2 last metabolites (*) cannot be calculated and the ratio between the ^12^C (sample metabolite) and ^13^C (standard) area was considered. For abbreviation see Figures 1B and 1D: G1P, glucose 1-phosphate; S7P, sedoheptulose 7-phosphate; Pentose5P, pentose 5-phosphate (ribose 5-phosphate, xylulose 5-phosphate and xylose 5-phosphate are undistinguished by IC-HRMS). (B) Enzymatic determination of the intracellular Gly3P concentrations in the parental (WT), ^*RNAi*^GK.ni and ^*RNAi*^GK.i cell lines grown in 10 mM glucose (blue), 10 mM glycerol (black) or both (grey). The absence of detectable amounts of Gly3P in cellular extracts from the ^*RNAi*^GK.i mutants maintained in glycerol (last column) is probably due to cell quiescence caused by the impossibility of this mutant to metabolize glycerol. The intracellular concentrations of metabolites are calculated with the assumption that the total cellular volume of 10^8^ cells is equal to 5.8 µl (Opperdoes et al., 1984). (C-D) Effect of increasing amounts of Gly3P (C) and F1,6BP (D) on HK activity determined on total extracts of the *T. brucei* PCF.

In conclusion, we describe here a new mechanism for the regulation of nutrient utilization based on the competition between two enzymes (kinases) for a common substrate (ATP). The sequestration of the two kinases in the glycosomes is key to this mechanism since these peroxisome-like organelles show limited or no nucleotide exchange with the cytosol at a metabolic timescale (Deramchia et al., 2014; Haanstra et al., 2014). Hence, the ATP pool available to glycosomal kinases is limited, offering a situation where significant excess of one kinase (here GK) can abolish almost completely the flux through the other one (HK). The competing kinases (HK and GK) catalyze the first step of their respective pathway and therefore control the utilization of their substrate (glucose and glycerol assimilation, respectively). The competition of given enzymes for a common substrate, in particular at branching metabolic points, is a well-known process used to finely tune metabolic fluxes (Siebert et al., 1969). However, as far as we know, this is the first example of an almost complete repression of one enzymatic activity (HK) by the large excess of another one (GK) competing for the same substrate, as a mechanism to control nutrient utilization. The consequence of this competition for one substrate, named “metabolic contest”, resembles the catabolic repression observed in prokaryotes and yeasts, albeit based on a completely different molecular mechanism, since no nutrient sensing and signaling are required. The advantage of the “metabolic contest” versus catabolic repression mechanism is mainly an immediate switch to the less preferred carbon source when the preferred one is exhausted and with a minimal energy cost since no regulation of gene expression (here HK or GK) is required.

## ACKNOWLEDGEMENTS

This work was supported by the Centre National de la Recherche Scientifique (CNRS), the Université de Bordeaux, the Agence Nationale de la Recherche (ANR) through grants ACETOTRYP of the ANR-BLANC-2010 call and GLYCONOV of the “Générique” call, the Laboratoire d’Excellence (LabEx) ParaFrap ANR-11-LABX-0024, the ParaMet PhD programme of Marie Curie Initial Training Network and the Institut Pasteur. Part of the work was carried out at MetaToul (Metabolomics & Fluxomics Facilities, Toulouse, France, www.metatoul.fr), which is part of the MetaboHUB-ANR-11-INBS-0010 national infrastructure (www.metabohub.fr). JCP is grateful to the INSERM for funding a temporary full-time researcher position.

## AUTHOR CONTRIBUTIONS

S.A., M.W., E.C., M.B., N.P., P.M, E.P., H.K.-B., C.A. and O.V. conducted the experiments; E.T., L.R., B.R., A.M., J.-C.P. and F.B. designed the experiments and wrote the paper.

## STAR METHODS

### Method Details

#### Trypanosomes and cell cultures

The PCF of *T. brucei* EATRO1125.T7T (TetR-HYG T7RNAPOL-NEO) were cultivated in glucose conditions at 27°C in the presence of 5% CO_2_ in SDM79 medium containing 10% (v/v) heat-inactivated fetal calf serum and 3.5 mg/ml hemin (Brun and Schonenberger, 1979). The glycerol-rich/glucose-free conditions were obtained by replacing glucose by glycerol in SDM79 and adding 50 mM N-acetyl-D-glucosamine that is a non-metabolized glucose analogue inhibiting glucose import (Azema et al., 2004), in order to prevent the consumption of the residual serum-derived glucose (final concentration in the medium: 0.5 mM) (Allmann et al., 2014). For the glycerol conditions, glycerol (10 mM) was added to the glucose/glycerol-depleted medium. The PCF of *T. congolense* TREU was cultivated at 27°C in the presence of 5% CO_2_ in TcPCF-3 medium composed of Eagle’s Minimum Essential Medium (Sigma-Aldrich) supplemented with 2.2 g/l NaHCO_3_, 25 mM HEPES, 0.1 mM hypoxanthine, 2 mM glutamine, 10 mM proline, 20% (v/v) heat-inactivated fetal calf serum and 3.5 mg/ml hemin (Coustou et al., 2010).

#### Inhibition of gene expression by RNAi

RNAi-mediated inhibition of gene expression of the *GK* genes was performed in the EATRO1125.T7T PCF by expression of stem-loop “sense-antisense” RNA molecules of the targeted sequences (Bringaud et al., 2000; Ngo et al., 1998) using the pLew100 expression vector, which contains the phleomycin resistance gene (kindly provided by E. Wirtz and G. Cross) (Wirtz et al., 1999). To do so, a 617-bp fragment of the *GK* gene (from position 460 to 1077) was introduced in the pLew100 vector to produce the pLew-GK-SAS plasmid. Briefly, a PCR-amplified 617-bp fragment using primers containing the antisense GK sequence with restriction sites added to the primers, was inserted into the HindIII and BamHI restriction sites of the pLew100 plasmid. The separate 615-bp PCR-amplified fragment using primers containing the sense GK sequence was then inserted upstream of the antisense sequence, using HindIII and XhoI restriction sites (XhoI was introduced at the 3′-extremity of the antisense PCR fragment). The resulting plasmid pLew-GK-SAS contains a sense and antisense version of the targeted gene fragment, separated by a 89-bp fragment, under the control of a PARP promoter linked to a prokaryotic tetracycline operator. The EATRO1125.T7T parental cell line was transfected with the NotI-linearized pLew-GK-SAS plasmid in 4-mm electroporation cuvettes with the Gene Pulser Xcell™ apparatus (Bio-Rad) by using as parameters 1,500 V, 25 µF, infinite resistance and two-pulse mode. Selection of the ^*RNAi*^GK mutant was performed in glucose-rich SDM79 medium containing hygromycin (25 µg/ml), neomycin (10 µg/ml) and phleomycin (5 µg/ml). Aliquots were frozen in liquid nitrogen to provide stocks of each line that had not been cultivated long term in medium. Induction of RNAi cell lines was performed by addition of 1 µg/ml tetracycline.

#### Western blot analyses

Total protein extracts of *T. brucei* PCF (5 × 10^6^ cells) were separated by SDS-PAGE (10%) and immunoblotted on TransBlot Turbo Midi-size PVDF Membranes (BioRad) (Harlow and Lane, 1988). Immunodetection was performed as described (Harlow and Lane, 1988; Sambrook et al., 1989) using as primary antibodies the rabbit anti-GK (1:5,000, gift from P. A. M. Michels, Edinburgh, UK) and the rabbit anti-PFR (1:10,000). Anti-rabbit IgG conjugated to horseradish peroxidase (BioRad, 1:5000 dilution) was used as secondary antibody. Revelation was performed using the Clarity Western ECL Substrate as described by the manufacturer (BioRad). Images were acquired and analyzed with the ImageQuant LAS 4000 luminescent image analyzer.

#### NMR spectroscopy experiments

3 × 10^7^ PCF of *T. brucei* were centrifuged at 1,400 g for 10 min, then the pellet was washed twice with PBS and the cells were incubated for 6 h at 27°C in 1.5 ml of incubation buffer (PBS supplemented with 5 g/l NaHCO_3_, pH 7.4) with 4 mM [U-^13^C]-glycerol and/or 4 mM glucose. This quantitative ^1^H-NMR approach was previously developed to distinguish between [^13^C]-enriched and non-enriched excreted molecules produced from [^13^C]-enriched and non-enriched carbon sources, respectively (Bringaud et al., 2015; Mazet et al., 2013; Millerioux et al., 2018). The viability of the cells during the incubation was checked by microscopic observation. At the end of the incubation, 500 µl supernatant were collected and 20 mM maleate were added in this aliquot as internal reference. ^1^H-NMR spectra were performed at 125.77 MHz on a Bruker DPX500 spectrometer equipped with a 5 mm broadband probe head. Measurements were recorded at 25°C with an ERETIC method. This method provides an electronically synthesized reference signal (Akoka et al., 1999). Acquisition conditions were as follows: 90° flip angle, 5000 Hz spectral width, 32 K memory size, and 9.3 sec total recycle time. Measurements were performed with 256 scans for a total time close to 40 min. Before each experiment, phase of ERETIC peak was precisely adjusted. Resonances of obtained spectra were integrated and results were expressed relative to ERETIC peak integration.

#### Mass spectrometry analyses of [^13^C]-incorporation into cellular metabolites

For analysis of [^13^C]-incorporation into intracellular metabolites, EATRO1125.T7T parental and mutant cell lines grown in SDM79 medium were washed twice with PBS and resuspended in an incubation solution (PBS containing either 2 mM [U-^13^C]-glycerol or 2 mM [U-^13^C]-glucose with or without the same amount of unlabeled glucose or glycerol). The cells were incubated for 2 h at 27°C before being collected on filters by fast filtration and prepared for MS analysis as described before (Ebikeme et al., 2010). Total sampling time was below 8 sec and the extraction of intracellular metabolites was carried out by transferring the filters containing the pellets into 5 ml of boiling water for 30 sec. The extracts were briefly vortexed (~2 sec), immediately filtered (0.2 µm) and chilled with liquid nitrogen. After freeze-drying, the dried extracts were resuspended in 200 µl Milli-Q water prior to analysis. Three replicates were taken from each culture, sampled and analyzed separately. The analyses of metabolites were carried out on liquid anion exchange chromatography Dionex™ ICS-5000+ Reagent-Free™ HPIC™ (Thermo Fisher Scientific™, Sunnyvale, CA, USA) system coupled to QExactive™ Plus high resolution mass spectrometer (Thermo Fisher Scientific™, Waltham, MA), as previously described (Millard et al., 2019). Central metabolites were separated within 48 min, using linear gradient elution of KOH applied to an IonPac AS11 column (250 x 2 mm, Dionex™) equipped with an AG11 guard column (50 x 2 mm, Dionex™) at a flow rate of 0.35 ml/min. The column and autosampler temperature were 30°C and 4°C respectively. Injected sample volume was 15 µl. Mass detection was carried out in a negative electrospray ionization (ESI) mode. The settings of the mass spectrometer were as follows: spray voltage 2.75 kV, capillary and desolvatation temperature were 325 and 380 °C respectively, maximum injection time 0.1 sec. Nitrogen was used as sheath gas (pressure 50 units) and auxiliary gas (pressure 5 units). The automatic gain control (AGC) was set at 1e6 for full scan mode with a mass resolution of 70,000. Identification of ^13^C carbon isotopologues distribution relied upon matching accurate masses from FTMS (mass tolerance of 5 ppm) and retention time using TraceFinder 3.2 software. To obtain ^13^C-labelling patterns (^13^C isotopologues), isotopic clusters were corrected for the natural abundance of isotopes of all elements and for isotopic purity of the tracer, using the in-house software IsoCor freely available at https://github.com/MetaSys-LISBP/IsoCor, and its documentation at https://isocor.readthedocs.io.

#### Determination of intracellular metabolite concentrations by IC-HRMS and enzymatic assays

For the quantification of intracellular metabolites, the *T. brucei* PCF EATRO1125.T7T cell line grown in glucose or glycerol conditions was sampled by fast filtration, as described above. The extraction of intracellular metabolites was performed as mentioned above before adding 200 µl of a uniformly [^13^C]-labeled *E. coli* cell extract as internal quantification standard (Wu et al., 2005). Then, metabolites were analyzed by IC-HRMS as described above with measured concentrations of metabolites expressed as a total cellular concentration assuming a volume of 10^8^ cells being equal to 5.8 µl (Opperdoes et al., 1984). The metabolites under examination include glucose 1-phosphate (G1P), 6-phosphogluconate (6PG), ribose 5-phosphate, ribulose 5-phosphate, xylulose 5-phosphate (considered together as pentose 5-phosphate, Pentose5P) and sedoheptulose 7-phosphate (S7P), in addition to those mentioned above.

For the enzymatic determination of Gly3P, the *T. brucei* PCF (5 × 10^7^ cells per sample) were washed in PBS and lysed in 100 µl of fresh 0.9 M perchloric acid. After centrifugation at 16,000 g at 4°C, 150 µl of H_2_O and 75 µl of the KOH (2M)/MOPS (0.5M) mix were added to the cellular pellet and incubated 5 min on ice. After centrifugation at 16,000 g at 4°C, the amounts of Gly3P contained in the supernatant was determined with the Amplite Fluorimetric G3P Assay kit, as described by the manufacturer (AAT Bioquest, Euromedex 13837 AAT).

#### Determination of glucose and glycerol consumption

To determine the rate of glucose and glycerol consumption, *T. brucei* PCF EATRO1125.T7T (inoculated at 10^7^ cells/ml) or *T. congolense* TREU (inoculated at 5 × 10^6^ cells/ml) were grown in 10 ml of SDM79 or TcPCF-3 media, respectively, containing 2.5 mM of glucose, 2.5 mM of glycerol or both (Brun and Schonenberger, 1979; Coustou et al., 2010). Aliquots of growth medium (500 µl) were collected periodically during the 10 h of incubation at 27°C. The quantity of glucose and glycerol present in the medium was determined using the “Glucose GOD-PAP” kit (Biolabo SA) and the “Glycerol assay kit” (Sigma-Aldrich), respectively. The amount of carbon source consumed at a given time of incubation (Tx) was calculated by subtracting the remaining amounts in the spent medium at Tx from the initial amounts at T0. Then, the rate of glucose and glycerol consumed per h and per mg of protein was calculated from the equation of the linear curve deduced from plotting carbon source consumption as a function of time of incubation. Importantly, we controlled that 100% of the cells remained alive and motile at the end of the 10 h of incubation.

#### Enzymatic activities

For enzymatic activities, PCF cells were washed in PBS (10’, RT, 900 g), resuspended in assay buffer and after addition of ‘Complete EDTA-Free’ protease-inhibitor cocktail (Roche), lysed by sonication (Bioruptor, Diagenode; high intensity, 5-10 cycles, 30sec/30sec on/off). Debris was spun down (15’, RT, 16,000 g) and the supernatants were used for protein determination with the Pierce BCA protein assay kit in a FLUOstar Omega plate reader at 660 nm. For higher throughput and smaller assay volumes all activity measurements were performed in a 96-well format with a FLUOstar Optima including an automated injection system. Malic enzyme activity was determined as quality control of the cellular extracts as described before (Allmann et al., 2013). The baseline reactions were measured for 2 min and the reactions were started by injection of the specific substrate or a combination of two (glucose, glycerol, malate, acetate) for each enzyme. The decrease/increase in absorbance at 350 nm was followed for 3-5 min. The rate was determined from the linear part of the progress curve and from this the specific activity was calculated. The buffer for determination of GK activity contains 100 mM triethanolamine pH 7.6, 2.5 mM MgSO_4_, 10 mM KCl, 0.6 mM ATP, 2 mM phosphoenolpyruvate, 0.6 mM NADH, ~1 U lactate dehydrogenase, ~1 U pyruvate kinase and 10 mM of glucose and/or glycerol (injected substrate). The buffer for HK measurements contains 100 mM triethanolamine pH 7.6, 10 mM MgCl_2_, 0.6 mM ATP, 0.6 mM NADP^+^, ~1 U glucose-6-phosphate dehydrogenase and 10 mM of glucose and/or glycerol (injected substrate) (Kralova et al., 2000). The same conditions are used for the determination of HK activity in the presence of acetate kinase, except the addition of 10 mM acetate and recombinant acetate kinase from *E. coli*. For the determination of HK activity the amount of 60 µg cellular protein was used per well, whereas for GK activity 6 µg per well were used.

