## Supplementary data for "“Metabolic contest”, a new way to control carbon source preference"

**Table S1 (Related to Figure 1).** Excreted end-products from metabolism of [U-^13^C]-glycerol and/or glucose by the parental (EATRO1125.T7T), *^RNAi^*GK.ni and *^RNAi^*GK.i procyclic *T. brucei* cell lines. The extracellular PBS medium of trypanosome incubated in the presence of 4 mM of one or two carbon sources was analyzed by ^1^H-NMR spectroscopy to detect and quantify excreted end-products.

| Cell line^a^ | Carbon source(s) metabolized^b^ |  |  | nmol/h/mg of protein | | | | | | | | | |
| --- | --- | --- | --- | --- | --- | --- | --- | --- | --- | --- | --- | --- | --- |
|  |  |  | n^c^ |  |  |  |  |  |  |  |  |  |  |
|  |  |  |  | Acetate | | Succinate | | Lactate | | Alanine | | TOTAL | |
| Parental (Glc) | **[U-^13^C]-Glycerol** |  | 9 | **1386** | **± 192.9** | **1123** | **± 205.3** | **ND**^d^ | | **ND** | | **2509** | **± 231.5** |
| Parental (Glc) | **[U-^13^C]-Glycerol** |  | 6 | **1445** | **± 209.4** | **911** | **± 101.3** | **ND** | | **ND** | | **2356** | **± 17.7** |
|  | Glucose |  |  | 25 | ± 26.6 | 17 | ± 24.2 | ND | | ND | | 42 | ± 25.3 |
| Parental (Glc) | Glucose |  | 6 | 1727 | ± 115.3 | 401 | ± 89.2 | 41 | ± 14.8 | 19 | ± 36.0 | 2188 | ± 144.6 |
| Parental (Glyc) | **[U-^13^C]-Glycerol** |  | 3 | **1357** | **± 193.2** | **1110** | **± 93.6** | **ND** | | **ND** | | **2468** | **± 143.2** |
| Parental (Glyc) | **[U-^13^C]-Glycerol** |  | 3 | **1336** | **± 126.1** | **890** | **± 170.2** | **ND** | | **ND** | | **2226** | **± 196.6** |
|  | Glucose |  |  | 89 | ± 44.2 | 49 | ± 16.3 | ND | | ND | | 139 | ± 46.8 |
| Parental (Glyc) | Glucose |  | 3 | 1413 | ± 142.0 | 690 | ±109.5 | 63 | ± 14 | 100 | ± 35 | 2266 | ± 151.8 |
| *^RNAi^*GK.ni | **[U-^13^C]-Glycerol** |  | 6 | **591** | **± 149.3** | **215** | **± 67.5** | **ND** | | **ND** | | **806** | **± 88.2** |
| *^RNAi^*GK.ni | **[U-^13^C]-Glycerol** |  | 6 | **429** | **± 97.6** | **155** | **± 44.2** | **ND** | | **ND** | | **584** | **± 129.3** |
|  | Glucose |  |  | 1341 | ± 63.3 | 736 | ± 216.6 | 60 | ± 10.4 | 3 | ± 11.3 | 2140 | ± 289.6 |
| *^RNAi^*GK.ni | Glucose |  | 6 | 1490 | ± 242.6 | 557 | ± 61.7 | 61 | ± 84 | 10 | ± 8.6 | 2118 | ± 257.1 |
| *^RNAi^*GK.i | **[U-^13^C]-Glycerol** |  | 6 | **35** | **± 14.4** | **8** | **± 6.2** |  | **ND** |  | **ND** | **43** | **± 20.5** |
| *^RNAi^*GK.i | **[U-^13^C]-Glycerol** |  | 6 | **59** | **± 5.4** | **8** | **± 0.6** |  | **ND** |  | **ND** | **66** | **± 10.4** |
|  | Glucose |  |  | 1608 | ± 215.7 | 696 | ± 160.3 | 78 | ± 8.5 | 3 | ± 11.3 | 2386 | ± 270.0 |
| *^RNAi^*GK.i | Glucose |  | 6 | 1531 | ± 349.1 | 656 | ± 109.0 | 75 | ± 8.7 | 16 | ± 3.1 | 2277 | ± 451.8 |

*^a^* The parental cells (EATRO1125.T7T) were cultivated several days in SDM79 containing 10 mM glucose (Glc) or in glucose-depleted SDM79 containing 10 mM glycerol and 50 mM GlcNAc (Glyc), before incubation in PBS. *^b^* Incubation conditions (carbon sources added to the PBS medium). *^c^* Number of duplicates. *^d^* Non detectable

**Figure S1 (related to Figure 1).** Growth curves of the parental (WT) and tetracycline-induced *^RNAi^*GK.i cell lines incubated in SDM79 medium containing 10 mM glucose (+ Glc, - Glyc), 10 mM glycerol (- Glc, + Glyc), both glucose and glycerol (+ Glc, + Glyc) or none of them (- Glc, - Glyc). Cells were maintained in the exponential growth phase (between 10^6^ and 10^7^ cells/ml), and cumulative cell numbers reflect normalization for dilution during cultivation.

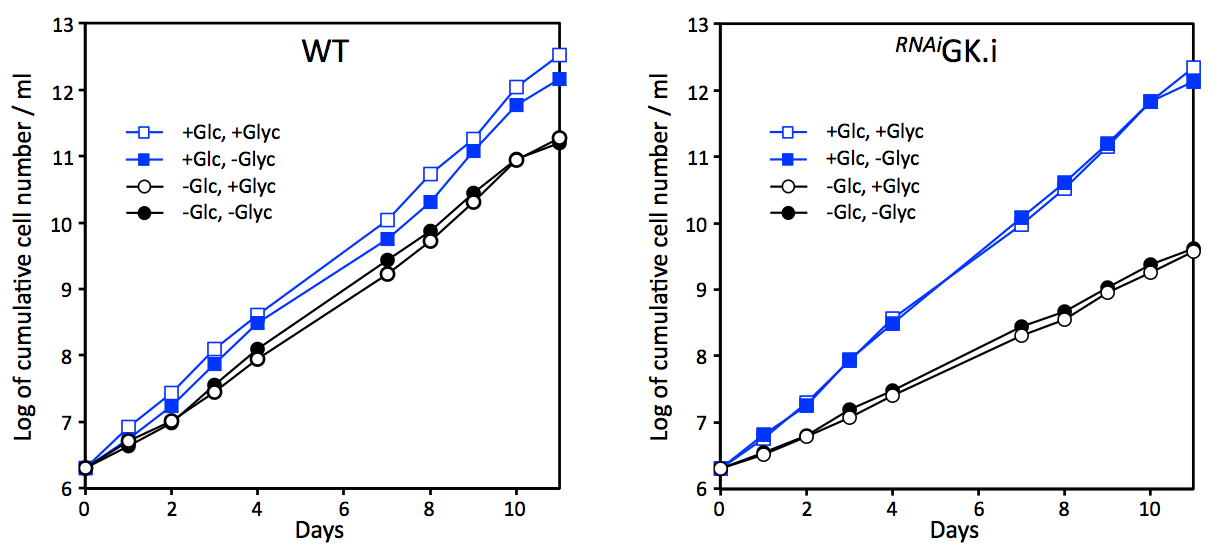

**Figure S2 (related to Figure 1).** Glycerol metabolism in other trypanosomatids. (A) GK and HK activities in total cell extracts of the procyclic form of *T. congolense* in the presence of glucose (Glc), glycerol (Glyc) or equal amounts of glucose and glycerol (Glc/Glyc). (B) Consumption of Glucose (left) and glycerol (right) by the *T. congolense* PCF incubated in glucose-rich (2 mM), glycerol-rich (2 mM) and glucose/glycerol (2 mM each) conditions. (C) Correlation between high GK/HK activity ratio and glycerol-preference. *^a^* HK activity in the presence of equimolar amounts of glucose and glycerol; *^b^* Ratio between the GK and HK activities; *^c^* Rate of glucose (Glc) or glycerol (Glyc) consumption; *^d^* Culture in the presence of 2 mM glucose (+Glc), 2 mM glycerol (+Glyc) or both (+Glc, +Glyc); *^e^* Data from Figure 1; *^f^* ND: not detectable; *^g^* Data from (B); *^h^* Data from (Pineda et al., 2018).

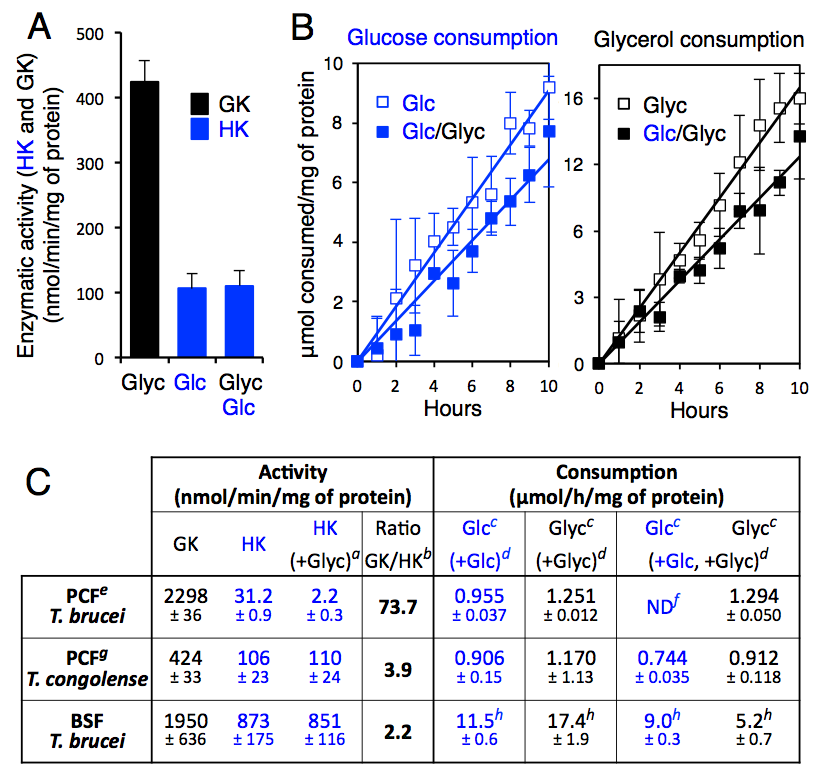
